# Coupling multiplex pre-amplification and droplet digital PCR for longitudinal monitoring of *ESR1* and *PIK3CA* mutations from plasma cell-free DNA

**DOI:** 10.1101/598847

**Authors:** Huilan Yao, Grant Wu, Subhasree Das, Crystal MacKenzie, Hua Gao, Victoria Rimkunas, Zhaojie Zhang, Stephanie Ferro, Amy Roden, Manav Korpal, Joanne Schindler, Peter G. Smith, Lihua Yu, Ping Zhu, Pavan Kumar

## Abstract

Here we report on the development of a sensitive and cost-effective method to longitudinally track *ESR1* and *PIK3CA* mutations from cfDNA in patients with metastatic breast cancer (MBC) using a streamlined and de-centralized workflow. Hotspot mutations in *ESR1* have been shown to cause resistance to aromatase inhibitor–based and anti-estrogenic therapies, while *PIK3CA* mutations have high prevalence in MBC. As a result, their utility as circulating biomarkers to predict or monitor response in the clinical development of investigational compounds has been the focus of many studies. Six regions in *ESR1* and *PIK3CA* genes containing 20 hotspot mutations were pre-amplified, followed by optimized singleplex ddPCR assays to detect allele frequencies of individual mutations. Without pre-amplification, the limit of detection (LOD) and limit of linearity (LOL) of individual ddPCR assays were at 0.05-0.1% and 0.25% level, respectively. With pre-amplification, the LOD and LOL were slightly elevated at 0.1-0.25% and 0.25-0.5% levels, respectively. High concordance was achieved to the BEAMing assay (Sysmex Inostics) for mutation positive assays (r=0.98, P<0.0001). In conclusion, coupling pre-amplification and ddPCR assays allowed us for the detection of up to 20 hot spot mutations in *ESR1* and *PIK3CA* with high sensitivity and reproducibility.

## Introduction

Breast cancer is the most common type of invasive cancer in women, which despite advances in detection and treatment, is responsible for over 500,000 deaths per year world-wide ^1^. Over 70% of primary breast cancers are positive for estrogen receptor α (ERα) and dependent on ERα signaling^2^. Anti-endocrine therapies suppress ERα signaling, either through estrogen deprivation (Luteinizing hormone-releasing hormone agonists, aromatase inhibition, (aromatase inhibitors, AI), or direct ER antagonism (tamoxifen or fulvestrant), and are widely employed in the clinic. Despite the initial effectiveness of anti-endocrine therapies, a vast majority of patients develop resistance. Although several mechanisms of resistance have been elucidated, it was only recently noted that recurrent mutations in *ESR1* are enriched in ~20-40% of patients with MBC treated with an AI ^3–10^. The fact that frequencies of *ESR1* mutations are extremely low in primary breast tumors and enriched in metastatic tumors, supports the potential functional role of these mutations in acquired resistance to anti-endocrine therapies. Indeed, subsequent preclinical studies confirmed the function of hotspot *ESR1* mutations in promoting resistance to various classes of endocrine therapies, perhaps by virtue of the constitutive activity gained by these mutants. The most frequently occurring *ESR1* mutations are clustered in the ligand-binding domain of ER^12,16,17^ between amino acid 534-538, though mutations at S463 and E380 are also described. *ESR1* mutations commonly co-occur with truncal PI3K mutations. In contrast to ER mutations, the frequency of *PIK3CA* mutations is unaffected by repeated AI exposure due to the truncal nature of these mutations ^4^. Up to 80% of *PIK3CA* mutations occur in hotspots within the helical (E542, E545, Q546) and kinase domains (M1043, H1047) of p110α ^11^.

Due to the prevalence of mutations in *ESR1* and *PIK3CA* in MBC, and their role in resistance to endocrine therapies, many clinical studies have characterized and monitored these mutations in cell free DNA (cfDNA) to explore their utility as circulating biomarkers to predict or monitor response to investigational compounds. Data from the FERGI trial showed that patients whose best response was complete response or partial response demonstrated robust decreases in *ESR1* and *PIK3CA* allele fraction after treatment with fulvestrant alone or in combination with pan-PIK3 inhibitor. In contrast, patients whose best response was stable disease or progressed disease displayed changes that were more variable ^12^. In a recent phase I trial of AZD9496 oral selective ER degrader, patients without *ESR1* ligand binding domain mutations (*ESR1*-LBDm) at baseline, or with a decline of mutations larger than 50% at C1D15, had a trend toward longer progression free survival (PFS), as compared to those who had less than 50% reduction of *ESR1*-LBDm cfDNA at C1D15 ^5^. Additionally, in the PALOMA-3 phase III study, longitudinal changes in *ESR1* and *PIK3CA* mutations in cfDNA were assessed for correlation with early efficacy of a CDK4/6 inhibitor. Early cfDNA dynamics in commonly occurring *PIK3CA* truncal driver mutations were shown to predict sensitivity to palbociclib. In contrast, subclonal mutations, such as *ESR1* mutations, were found to be weak predictors of outcome ^4^. Although the dynamics of mutations may vary due to different treatments, *ESR1* and *PIK3CA* mutations in cfDNA remain interesting biomarkers for future clinical studies.

Over the last decade, cfDNA has become an attractive and acceptable sample type for monitoring mutation changes ^13–15^. It has many advantages compared to tissue biopsy characterization. First, detection of mutations in blood samples represents a holistic mutation assessment, while a tissue biopsy performed at a specific location inevitably has sampling bias due to the heterogeneous nature of the tumor in addition to absence of any tumor characterization from other metastatic sites. Moreover, obtaining cfDNA from a blood draw is a minimally invasive procedure and can allow for longitudinal tracking throughout treatment. In contrast, obtaining multiple tissue biopsies from patients with advanced disease is typically not feasible. And finally, the short half-life of cfDNA in circulation, which is <2 hours ^16^, provides a “real-time” snapshot of disease burden. Many studies monitoring patients during treatment have shown that the tumor mutations in cfDNA (ctDNA) dynamics correlate with treatment response in MBC ^17^, non-small cell lung cancer (NSCLC) ^18^, colon cancer ^19,20^, melanoma ^21,22^, and may identify response or progression at an earlier time point than radiologic detection ^18–20,22^.

Detection of tumor mutations in cfDNA can be accomplished using various technologies. The first one is quantitative PCR (qPCR), which was first used to detected *KRAS* mutations in 1994 ^23^. Many qPCR platforms with regulatory approval are now on the market, such as Cobas (Roche), Therascreen (Qiagen), AmyDx Super ARMs qPCR system, and Idylla (Biocartis) ^24^. The second category is digital PCR, which was introduced in 1999 ^25^. It enabled the absolute quantification of rare mutant fragments and represented the next step in the evolution of qPCR. A modified version of this technique using beads in emulsions and flow cytometry, now known as BEAMing technology (Sysmex Inostics), is currently in use for detection of several mutations including *ESR1*, *PIK3CA* and so on ^12,26^. More recently, droplet digital PCR (ddPCR) on the Bio-Rad platform is expanding its footprint from the research setting into the clinical testing space. Not only the platform obtained 510k approval by FDA, but also the first ddPCR based assay for monitoring and quantifying residual CML disease was approved. The third category is next generation sequencing (NGS) based technologies. In contrast to PCR based approaches and platforms that focus on hot spot mutations in specific genes, the application of NGS based technologies in cfDNA surfaced in 2012 with a panel of tagged amplicons (Tam-Seq) for advanced ovarian cancer ^27^. This method was later applied to monitoring mutations in MBC ^17^. Since then, many different targeted sequencing approaches have been developed, such as Enhanced Tam-Seq ^28^, Safe-Seq ^29^, CAPP-Seq ^30^, and Digital sequencing ^31,32^ and are the basis of several commercially available panels, such as Guradant360 ^31^, FoundationACT ^33^ and Archer Reveal ctDNA panel ^34^ which have gradually expanded the gene content of the panels while still maintaining sensitivities typically required for cfDNA application.

When it comes to mutation monitoring, each technology has its own pros and cons. While NGS panel can simultaneously detect multiple mutations, it is hard to reliably detect mutations that are less than 1% ^35^. Stetson and colleagues did a replicate study across multiple ctDNA sequencing vendors and found a high degree of variability among assays particularly at low allele fractions (AF<1%) ^36^. Additionally, intra-assay variability was high at low frequency mutations ^37^. The BEAMing assay has become widely used in clinical trials, as it has demonstrated the highest reported sensitivity amongst digital PCR assays (down to 0.05% for some assays) while also allowing for detection of 22 mutations in the *ESR1*, PI3KCA and AKT genes due to its multiplexing nature. However, this assay as well as other high complex NGS assays necessitates centralized laboratory testing due to high requirement of both technical expertise and specialized analysis pipelines. These platforms do not allow for cost-effective, fast turn round testing at local labs. In contrast, droplet digital PCR has potential to be adopted in local labs with more streamlined workflows. Singleplex ddPCR assays, while very sensitive, require a significant amount of input material to detect each mutation. Therefore, usually 4-5 mutations are measured per sample. To circumvent some of the abovementioned limitations with existing analytical technologies, we herein describe a new method that couples multiplex pre-amplification and droplet digital PCR method for detecting and tracking up to 20 *ESR1* and *PIK3CA* mutations using a small amount of plasma cfDNA. It provides a sensitive, cost-effective, de-centralized and customized solution for longitudinally tracking mutations and can be customized by individual labs for their own mutations of interest.

## Result

### Panel design and Assay workflow

To design a panel to track *ESR1* and *PIK3CA* mutations, published papers for mutation selection in MBC patients were analyzed (Supplementary Table 1). Twenty-one mutations were selected (Table 1), as by combining all mutations, we could track more than 55% of the MBC patient population (Supplementary Table 2). Amongst all the published ddPCR panels (Supplementary Table 1), the OncoBEAM CLIA panel 1 (Sysmex Inostics) ^12^, containing both *ESR1* and *PIK3CA* mutations, was found to be most comprehensive and served as our reference assay: not only for content, but also as a benchmark for accuracy and internal validations.

**Table 1.** Limit of Detection (LOD) and Limit of Linearity (LOL) of ddPCR assays before and after pre-amplification.

| Gene | AA position | cDNA Change | Tm | Without pre-amplification |  | With Pre-amplification |  |
| --- | --- | --- | --- | --- | --- | --- | --- |
|  |  |  |  | LOD | LOL | LOD | LOL |
| ESR1 | V534E | c.1601T>A | 56 | 0.05% | 0.05% | 0.05% | 0.25% |
| ESR1 | P535H | c.1604C>A | 56 | 0.10% | 0.25% | 0.10% | 0.25% |
| ESR1 | L536H | c.1607T>A | 56 | 0.05% | 0.05% | 0.05% | 0.05% |
| ESR1 | L536R | c.1607T>G | 56 | 0.05% | 0.25% | 0.10% | 0.25% |
| ESR1 | L536Q | c.1607_1608TC>AG | 56 | 0.05% | 0.25% | 0.25% | 0.25% |
| ESR1 | Y537S | c.1610A>C | 56 | 0.05% | 0.05% | 0.10% | 0.10% |
| ESR1 | Y537N | c.1609T>A | 56 | 0.05% | 0.10% | 0.05% | 0.10% |
| ESR1 | Y537C | c.1610A>G | 56 | 0.10% | 0.25% | 0.50% | 0.50% |
| ESR1 | D538G | c.1613A>G | 56 | 0.05% | 0.05% | 0.25% | 0.25% |
| ESR1 | S463P | c.1387T>C | 56 | 0.10% | 0.25% | 0.25% | 0.25% |
| ESR1 | E380Q | c.1138G>C | 56 | 0.05% | 0.10% | 0.25% | 0.25% |
| PIK3CA | C420R | c.1258T>C | 56(53-56) | 0.10% | 0.25% | 0.25% | 0.50% |
| PIK3CA | E542K | c.1624G>A | 55 | 0.05% | 0.25% | 0.05% | 0.25% |
| PIK3CA | E545G | c.1634A>G | 55(55-53) | 0.25% | 0.25% | 0.25% | 0.25% |
| PIK3CA | E545K | c.1633G>A | 55 | 0.10% | 0.10% | 0.10% | 0.25% |
| PIK3CA | Q546K | c.1636C>A | 55(55-53) | 0.05% | 0.05% | 0.05% | 0.05% |
| PIK3CA | M1043I | c.3129G>A | 53 | 0.10% | 0.25% | 0.10% | 0.25% |
| PIK3CA | M1043I | c.3129G>T | 53 | 0.10% | 0.10% | 0.25% | 0.25% |
| PIK3CA | H1047L | c.3140A>T | 56 | 0.25% | 0.25% | 0.25% | 0.25% |
| PIK3CA | H1047R | c.3140A>G | 56 | 0.10% | 0.25% | 0.10% | 0.25% |
| PIK3CA | H1047Y | c.3139C>T | 55 | 0.25% | 0.25% | 0.25% | 0.25% |

A summary of our internal assay workflow is outlined in Figure 1. In brief, cfDNA was extracted from 2-3 mL of plasma and quantified using the LINE-1 qPCR assay to determine amplifiable copies or number of genome equivalents. 7.5ul of extracted cfDNA was subjected to multiplex PCR pre-amplification using gene specific primers to result in 6 amplicons covering all hot spot mutations of interest (Table 1). The input amount of pre-amplified material for ddPCR reactions was then optimized to maximize the possibility of having a single target DNA molecule per droplet. Pooled post pre-amplification product was diluted 1:10^6^ and used to quantify total copies of each pre-amplified amplicon using off the shelf ddPCR assays (BioRad). Appropriate sample level dilutions were then prepared to ensure 30,000-60,000 copies of each corresponding amplicon went into the final ddPCR reaction to enable a theoretical LOD of 0.01%. The final allele burden quantification was performed as a singleplex reaction in triplicate on the BioRad ddPCR platform for each relevant mutation with appropriate positive (a 3-5-point standard curve made with wildtype/mutant synthetic gBlocks) and negative (WT gBlock and water) controls.

**Figure 1.**
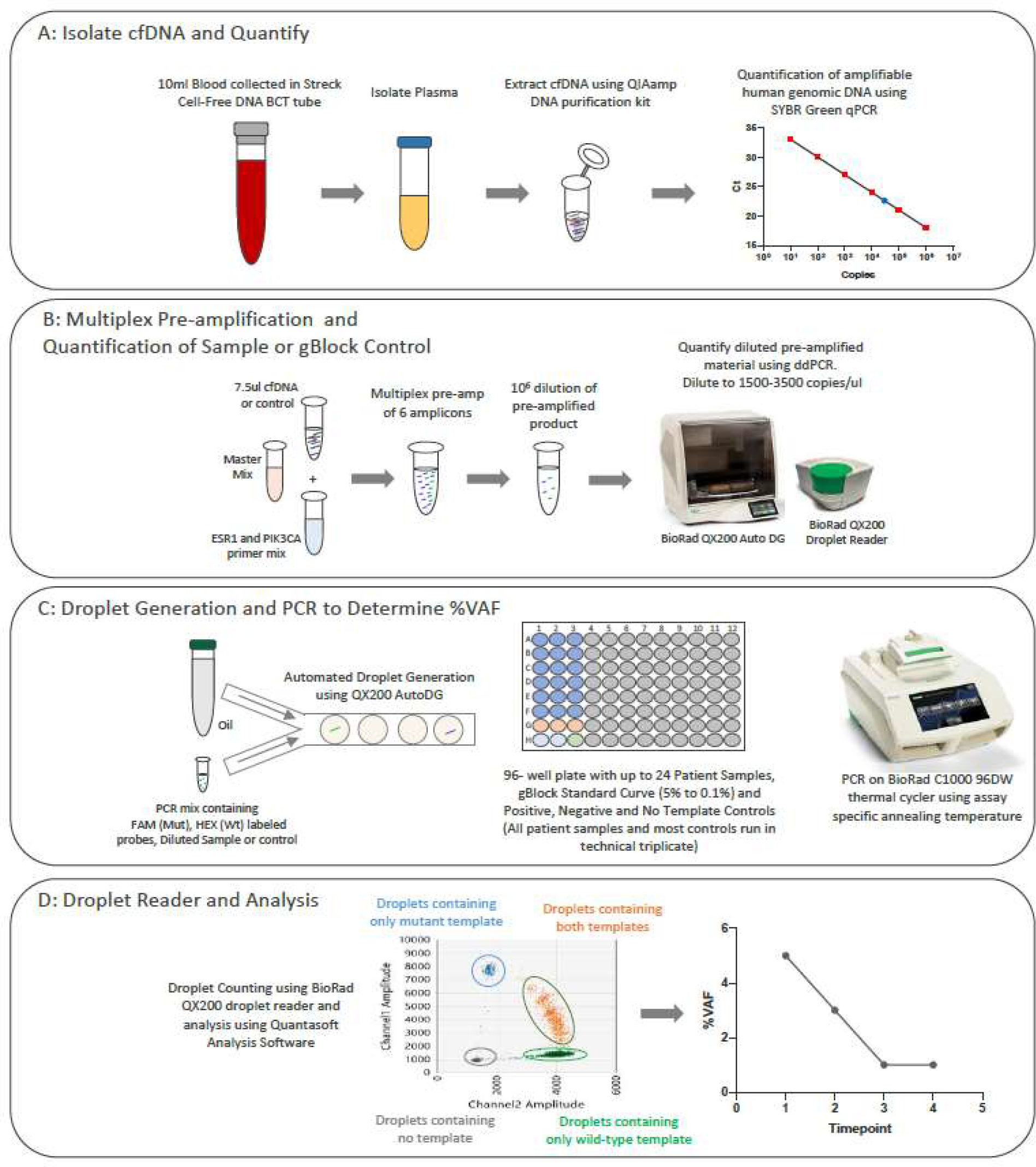
Workflow of the in-house ddPCR assays. (A): Peripheral blood is collected from patient into 10 mL Streck Cell-Free DNA BCT tube and plasma is isolated. cfDNA is extracted from 2-3 mL of plasma using Qiagen’s QIAamp DNA Purification kit. The LINE-1 SYRB Green qPCR assay is used to quantify amount of amplifiable genomic DNA and results are reported as Genomic Equivalents (GE’s). (B): 7.5μL of cfDNA or gBlock control is used for multiplex pre-amplification (16 cycles followed by 15 cycles of touch down PCR) of 6 amplicons targeting frequently mutated hotspot regions in *ESR1* and *PIK3CA* genes. Pre-amplification product is diluted to target 1500 to 3500 copies/μL and is subsequently quantified with a preliminary round of ddPCR using off the shelf ddPCR assays to accurately quantify total copies of each pre-amplified amplicon. (C): Pre-Amplified samples are appropriately diluted to target one DNA molecule per droplet. Droplets are generated for each patient sample and gBlock or water control using the Bio-Rad Auto Droplet Generator (DG). Each patient sample is run in technical triplicate alongside a gBlock standard curve (% mutant gBlock ranging from 5 to 0.1%) in a 96 well plate. PCR amplification is done using assay specific anneal temperature on the BioRad C1000 thermal cycler with 96 deep well block. (D): QuantaSoft Analysis Software was used for analysis. The mutant positive droplets (FAM) containing amplified product were differentiated from negative or wild-type (HEX) droplets by applying a threshold using 2D Amplitude data. Total number of positive droplets was used to calculate % variant allelic frequency (VAF) for each patient sample.

### Multiplex PCR Design and Optimization

To cover the mutations shown in this panel, six amplicons (Supplementary Table 3) were designed to amplify targeted regions of the *ESR1* and *PIK3CA* gene. Primers for each amplicon were tested individually for amplification yield and specificity using standard PCR followed by Sanger sequencing (Supplementary Figure 1). To minimize non-specific priming and polymerase enzyme induced error, a hot start, high-fidelity proofreading enzyme was used. A touchdown step was added before the main amplification cycle to ensure PCR specificity (see material and methods).

To ensure balanced amplification of each amplicon when performing multiplex PCR (MX-PCR), primer concentration for each of the targets, as well as number of amplification cycles were optimized. The ratio of the highest amplicon yield to the lowest amplicon yield (balance ratio) was used to assess the balance of amplification. Prior to the MX-PCR, amplification yield of each amplicon was measured individually using standard PCR coupled with gel electrophoresis. It was observed that amplicons P420 and P1047 had the lowest amplicon yield amongst all six targets. Three primer pools (A-C in Supplementary Table 4) were tested to determine the individual primer concentration required to achieve balanced amplification efficiency across all targets. Primer set A had equal amounts of all primers while primer sets B & C had higher primer concentrations for P420 and P1047 to address the lower amplification efficiencies of these primer pairs observed in the singleplex reactions. Coverage analysis was performed using an Ion Torrent PGM NGS system (data not shown).

Based on initial NGS results, additional primer pools were tested (D-I in Supplementary Table 4) with varying concentrations of the amplicon specific primers to further normalize the amplification efficiency of all amplicons. Coverage analysis identified Ratio D-H resulting in relatively balanced amplification, with balance ratios of <5 (Figure 2A). As expected, 15 cycles of pre-amplification resulted in slightly increased balance ratios compared to 10 cycles due to further magnification of differences in amplification efficiencies among different primer pairs with increasing pre-amplification cycles. Given that the differences between 10 cycle and 15 cycle amplification were small, 15-cycle pre-amplification was chosen for subsequent assay validation to ensure enough material got pre-amplified with the lowest sample input amount. Ratio E achieved the lowest balance ratio and was selected for further testing. To determine the reproducibility of the multiplex reaction, different input amounts ranging from 5 - 150 ng of K562 cell line gDNA samples were pre-amplified in four independent experiments and submitted for NGS coverage analysis. As shown in Figure 2B, balance ratios remained <5, with CVs <15% for all the input amounts tested.

**Figure 2.**
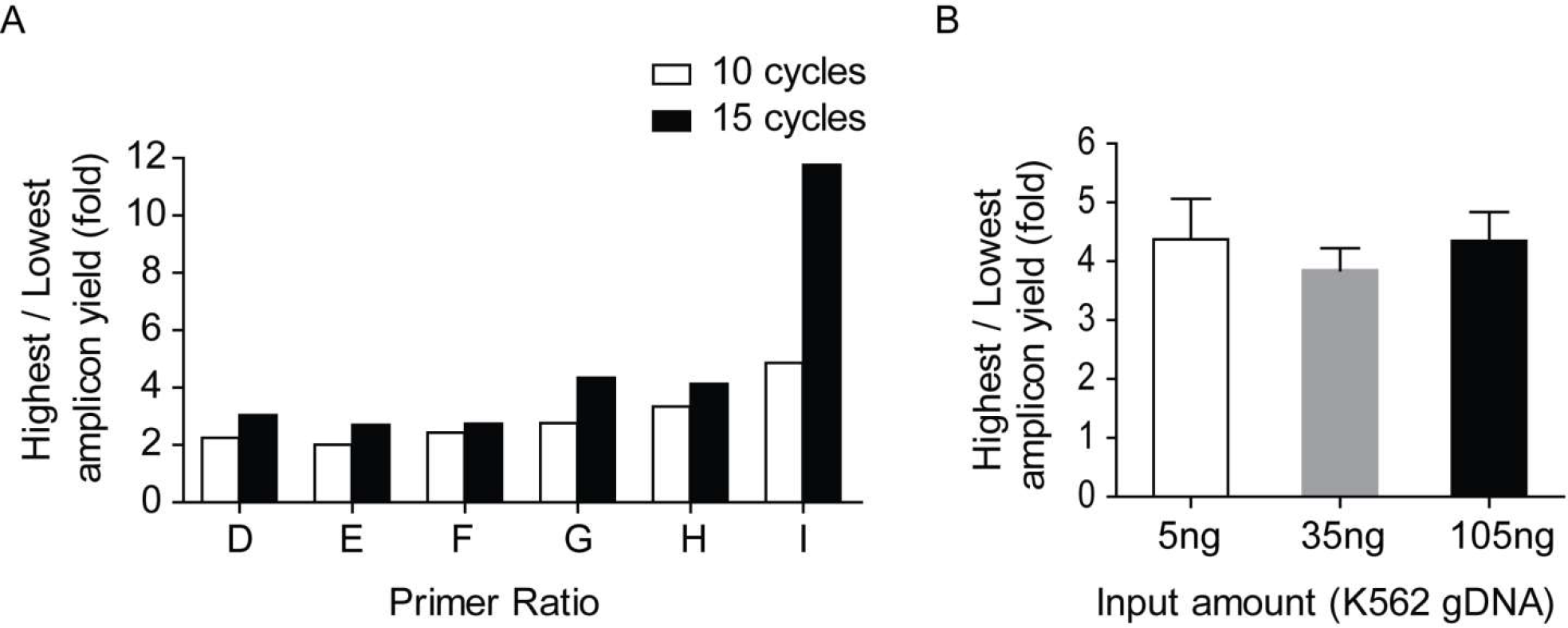
Optimization of multiplex-PCR condition. (A) Coverage analysis of the MX-PCR with various primer ratios and differing main amplification cycle number. The Y axis shows the ratio of the highest amplicon yield to the lowest amplicon yield measured by NGS, which is used to measure the evenness of pre-amplification of six amplicons. The X axis shows different primer pools with different ratios of individual primer sets. For detail information, see Supplementary Table 3. (B) Reproducibility of multiplex PCR condition using different amount of input DNA. Primer set pool E and 15 cycles for main amplification were used for pre-amplification. Mean and standard deviation of four replicates are shown here.

### Specificity assessment and optimization

Given that patients could have polyclonal mutations on the same nucleotide position or adjacent positions, cross-reactivity between individual assays had to be monitored and minimized. To assess the cross-reactivity, a 500 bp long mutant gBlock flanking the mutation of interest was designed for each individual mutation and used as positive control. A 1286 bp long WT gBlock which contains wild type sequences from all the six amplicons served as a positive control for wild type nucleotides. Nine assays that detect mutations clustered around *ESR1* amino acids 534-538 were selected from the panel and their cross-reactivity to the rest of the eight gBlocks was determined. This 9 X 9 assay matrix revealed cross-reactivity between assays that detected the mutations at the same nucleotide position, but not mutations at the adjacent nucleotide positions. For example, initial testing revealed cross-reactivity between two *ESR1* assays, Y537S, (c.1610A>C) and Y537C (c.1610A>G) (Figure 3A). The Y537S assay should only detect A>C while the Y537C assay should only read out A>G mutation at the c.1610 position. But at the standard annealing temperature of 55°C, the Y537S assay resulted in signal (positive droplets) from the c.1610G template (Figure 3A-b). No cross reactivity was observed when specific assays were targeting different nucleotide position changes (even separated by a single nt). For example, neither the Y537S assay (c.1610A>C) nor Y537C assay (c.1610A>G) detected positive signal from the control gBlocks for Y537N mutation (c.1609T>A) (Figure 3A-d, h). To mitigate this cross-reactivity, annealing temperatures (Tm) were optimized individually across all assays. Increasing Tm from 55°C to 56°C reduced the cross talk of assay Y537S to c.1610G sequence, but did not affect the detection of the correct c.1610C template (Figure. 3B). The optimized Tm for each individual assay is shown in Table 1.

**Figure 3.**
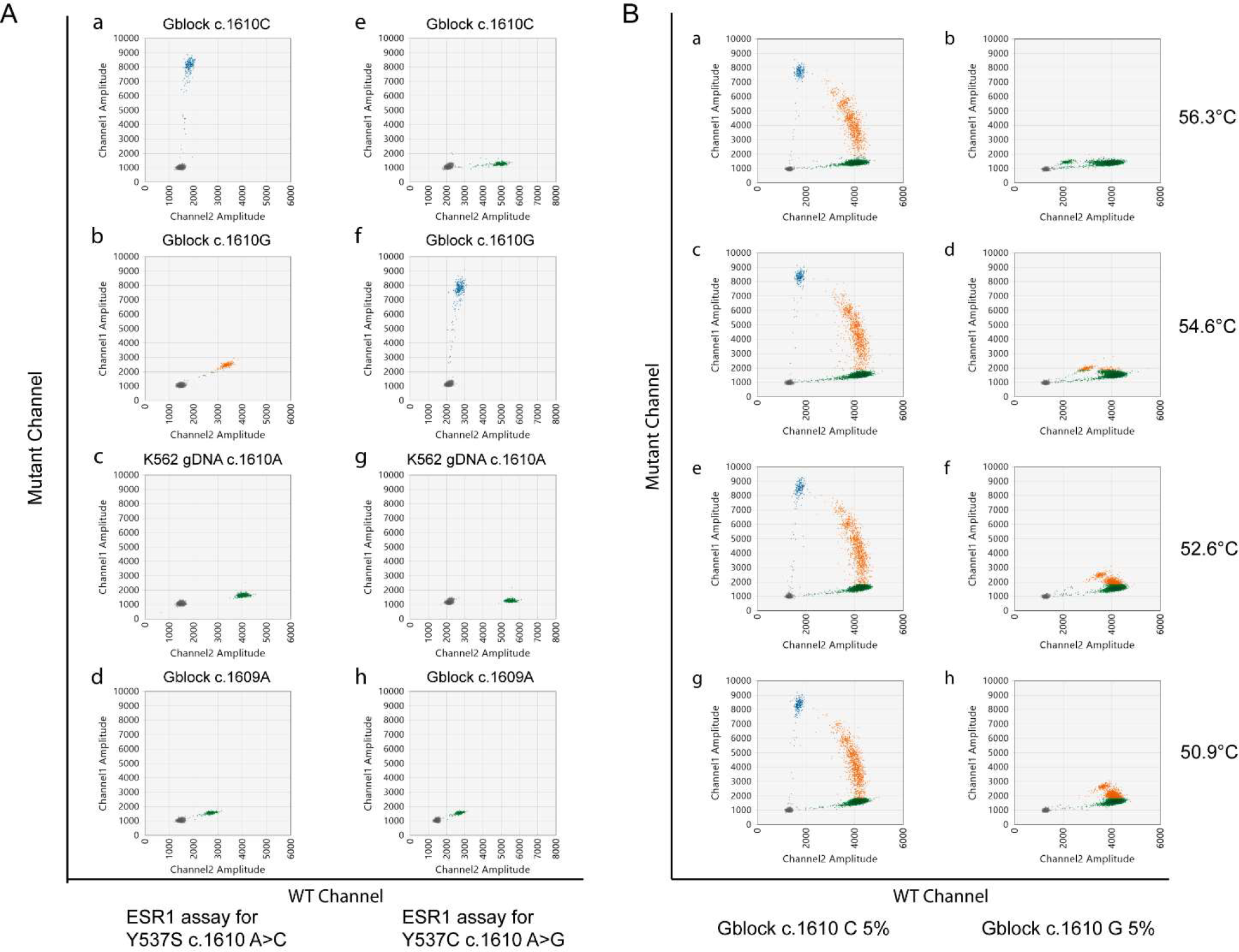
Optimization of Tm to minimize crosstalk between assays that detect nucleotide changes at the same position. For both (A) and (B), 2-dimentional amplitude of ddPCR assays are shown. The x-axis shows the signal amplitude from the wildtype channel, and the y-axis shows the signal amplitude from the mutant channel. Threshold were set up so the droplets contains only positive signal in wildtype channel are green, droplets contains only positive signal in mutant channel are blue, and droplets contain both wildtype and mutant signals are orange. Grey droplets are empty droplets that do not contain either wildtype signal or mutant signal. (A) Examples of crosstalk between assays that detect nucleotide changes at the same position, but not adjacent position at a suboptimal Tm. The top panels (a-d) were measured with assay *ESR1* Y573S, while the bottom panels (e-h) were measured with assay *ESR1* Y537C. Each panels from top to bottom contains specific gBlocks that is C at ESR1 c.1610 position, G at *ESR1* c.1610, K562 gDNA which is A at the *ESR1* c.1610, and A at c.1609 position. Note at the suboptimal Tm, mutant assay that bind to the c.1610C could crosstalk to gBlocks that has c.1610G (b), resulting in double positive droplets that would have led to false positive signal. But both assays do not crosstalk to gBlocks that has c.1609A (d, h). (B) Optimization of Tm to minimize crosstalk between assays. The left column (a,c,e,g) 4 samples are 5% gBlock ESR1 c.1610C while the right column (b,d,f,h) 4 samples are 5% gBlock ESR1 c.1610G. All samples were measured with assay *ESR1* Y573S (that should detect c.1610C) at different annealing temperatures. Note at Tm 56°C, mutant assay that bind to the c.1016C does not crosstalk to gBlocks that has c.1016G (a, b).

### Assay validation using contrived samples

One concern of pre-amplification was to lose the linear quantification property of typical ddPCR assays due to uneven or inconsistent pre-amplification across amplicons and/or individual samples. To this end, 5-point standard curves with gBlocks at mutant allele burden of 10%, 5%, 1%, 0.5%, 0.25%, 0.1%, 0.05% and 0% were used to determine the LOD and LOL for each assay with or without pre-amplification. Without pre-amplification, the majority of the assays had a LOD at the 0.05-0.1% level, with 0.05% being the last point on the standard curve. However, for the majority of assays the LOL was typically higher at 0.25%. So, while we could detect mutant allele burden of 0.25%, 0.1% and in some cases (depending on the assay) 0.05%, we lost the linearity at these lower levels and the assay became a qualitative readout. Not surprisingly, after pre-amplification, LOD increased to 0.25%-0.5% for 7 assays, however, LOL was usually maintained at 0.25% level (Table 1 and Figure 4). A few assays had a slightly higher LOL at 0.5% level, potentially due to increase in background signals from the WT gBlocks after pre-amplification which required a higher and more stringent threshold cut-off to eliminate false positive signals, thus leading to increased LOL.

**Figure 4.**
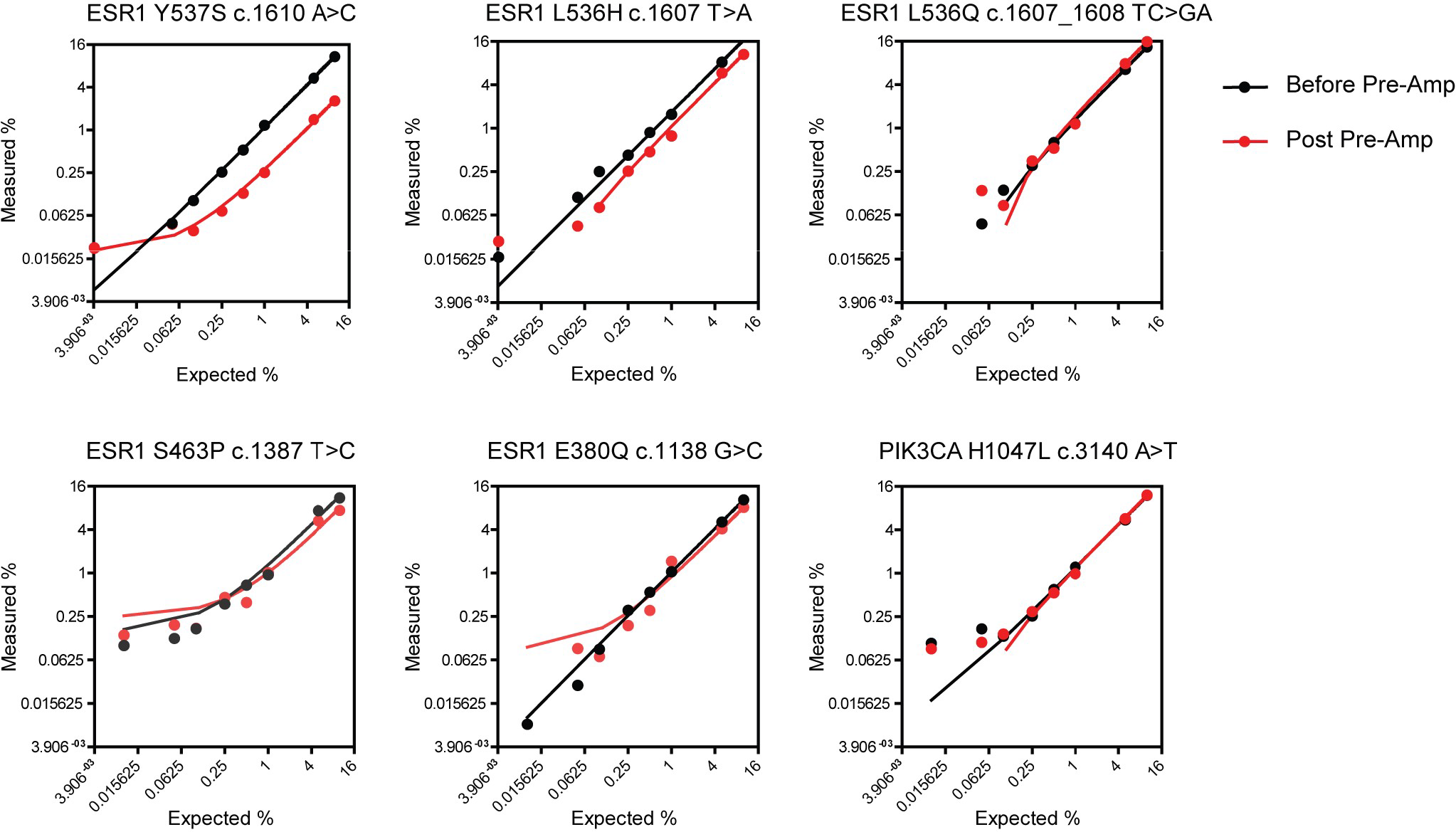
Determine the limit of detection and limit of linearity of ddPCR assays. Six representative assays were shown as indicated on top of each panel. Standard curves were prepared with each mutant *ESR1* gBlocks and wildtype gBlocks to generate a standard curve representing mutant allele burden of 10%, 5%, 1%, 0.5%, 0.25%, 0.1%, 0.05% and 0% (WT gBlock). Average measured VAF (%) of triplicates are shown before pre-amplification (black dots) and after pre-amplification (red dots). The lines on the graph are linear regression for each data set. The last dot in the linear regression line determined the limit of linearity of the assay; the last dot above the 0% sample determines the limit of detection of the assay.

We wanted to further understand and characterize the background sample type that was used to define the LOD of the assay. The WT gBlock used, although being reproducible and quantifiable, also represented a very high concentration of a single WT amplicon differing from the mutant target by a single nucleotide as opposed to a more complex real-life background type that would be randomly fragmented gDNA. The first alternate background sample tested for this purpose was a cfDNA reference standard set from Horizon. This was cell line DNA fragmented to average lengths of 160bp, that more closely resembled cfDNA extracted from plasma. The fragmented cell line was PIK3CA E545K mutant positive, thereby also serving as a negative control to assess the background for all the ESR1 assays. Background for mutations from these commercial cfDNA samples was either comparable to or lower than the WT gBlock samples for many assays (Figure 5). This trend was further confirmed when we assessed cfDNA from patient samples known to be WT for the mutations in question providing further confidence that the LOD set using the WT gBlock was stringent and conservative.

**Figure 5.**
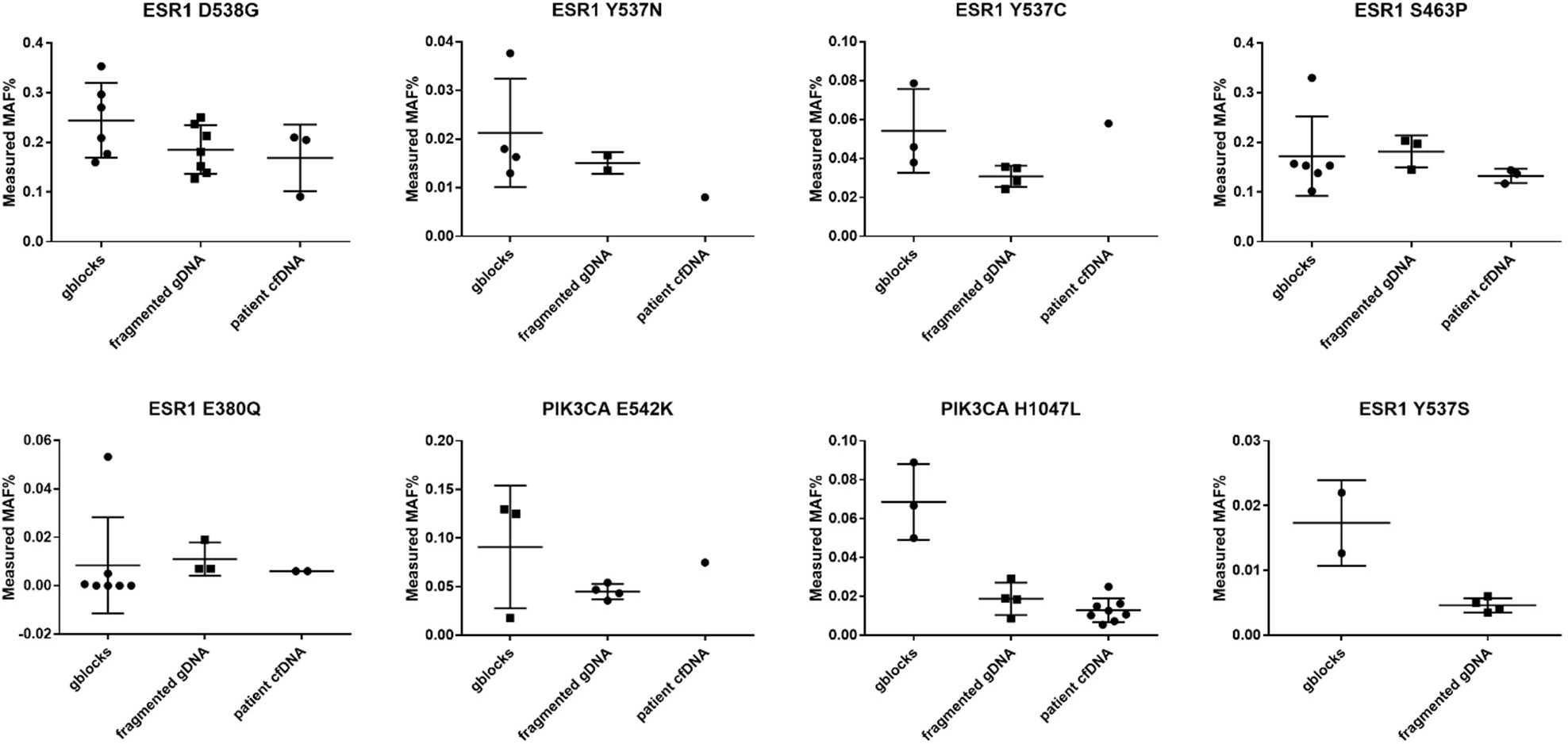
Background comparison among wildtype gblocks, fragmented gDNA, and patient plasma cfDNA. All samples used were supposed to be negative for mutations; therefore, the measured Mutant allele frequencies (MAF) % reflects the background using each of the sample types. Assay names are shown on top of each panel. Representative assays are shown there.

Assay reproducibility was assessed using the dilution standard curves made using the gBlocks and were run in triplicate (Supplementary Table 5). As expected, there was more variation observed (30-40% CV) at the lower end of the dilution curve (0.1-0.2%) as compared to <20% CV when tracking allele burden of >1%.

### Assay performance in clinical samples and impact of different amount of input material

To evaluate assay performance in our target patient population, we utilized 90 patient samples from the ongoing phase I clinical trial (NCT03250676) in ER+/HER2-MBC. Among them, 38 samples have been tested using both Inostics BEAMing assays ^12^ and the in-house ddPCR assays described here. The first question was to understand the input cfDNA quantity and quality and how it affected allele burden quantification. While several previous studies ^10,38^ used total amount (ng) of cfDNA as a measurement of input; for ddPCR assay, it is important to know how many copies of genomic equivalence (GE) were used as input which then sets the sample driven LOD. For example, if only 50 genomes worth of cfDNA was used as input, then even with the most sensitive assay that would detect a single mutant genome out of 50 genomes, the theoretical LOD would be 1/50 or 2%. So even though, the technical LOD based on contrived samples would be 0.25% (for e.g.) the amount of input cfDNA would be a limiting factor. To accurately assess the number of GEs in the cfDNA sample, the LINE-1 real time qPCR assay was used ^13^. To verify that the concentration measured by LINE-1 represented the concentration of the six regions from *ESR1* and *PIK3CA*, the copy number of each amplicon was measured using one representative assay from that region using our custom ddPCR assays. As shown in Figure. 6A, the LINE-1 assay was a good surrogate for concentrations from other regions. Representative analysis from 2 patients is shown where GEs from LINE-1 were within 50% of the average copies of the multiple amplicons being assessed.

**Figure 6.**
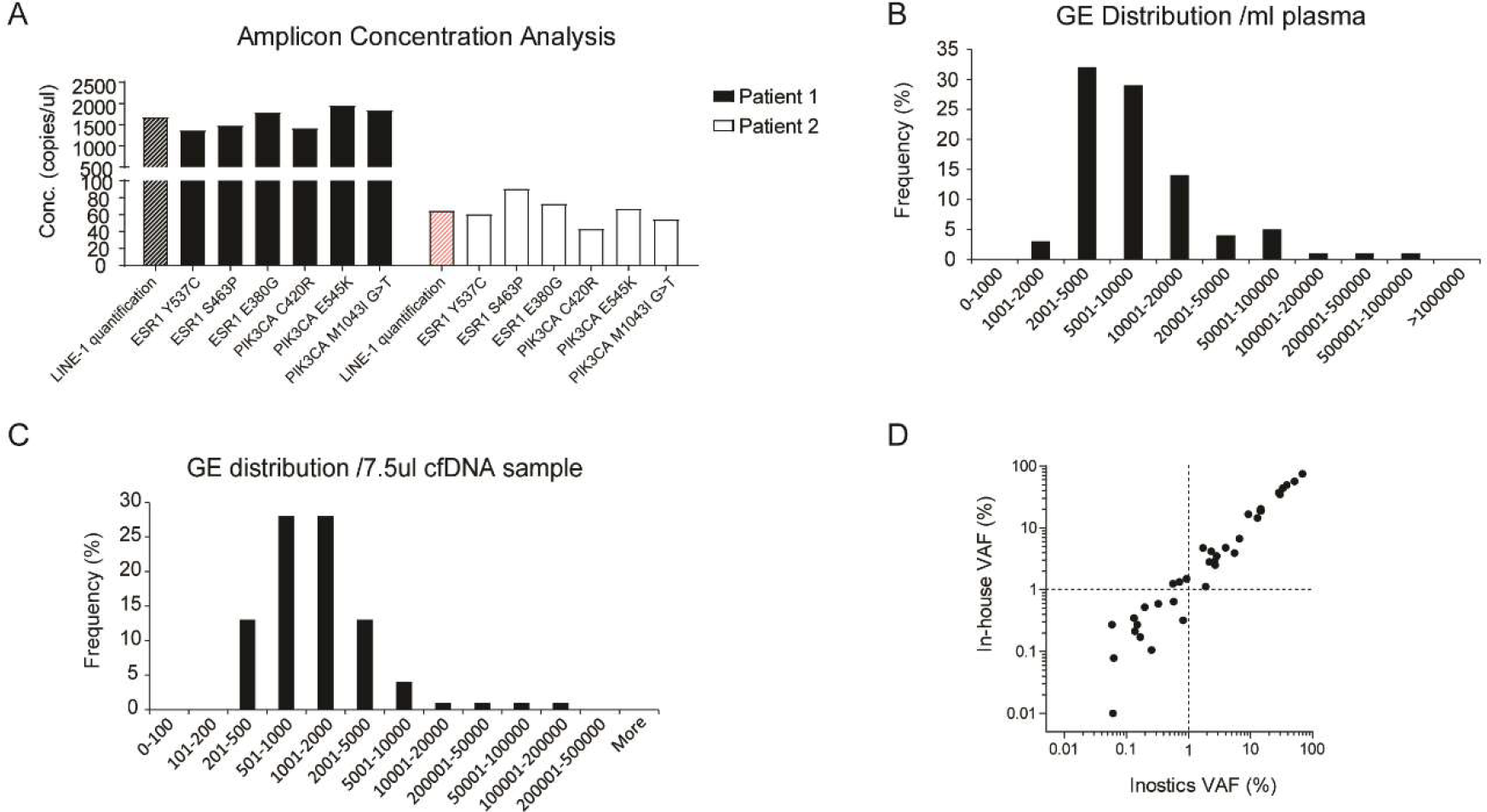
Performance of ddPCR assays in patient cfDNA samples. (A) Amplicon concentration analysis of two patients samples using LINE-1 realtime assay or ddPCR assays for individual amplicons. All assays were done without pre-amplification. (B) Histogram of cfDNA concentration of 90 patients’ cfDNA samples. cfDNA concentrations were shown as genomic equivalence/mL of plasma, measured by LINE-1 real time PCR assay. (C) Histogram of cfDNA input quantity for pre-amplification. The GE number in 7.5ul cfDNA is shown. For (B) and (C), x-axis shows the bin ranges. Y-axis shows the frequencies of samples that fall into each bin. (D) Correlation between VAF (%) measured by Inostics assay and in-house ddPCR assays.

Next, we determined if the current assay LOD was compatible with sample derived LOD. The range of GEs from 2 mL of patient samples was 1,443-980,577 copies/mL plasma; with the median GE of cfDNA being 6,327.75 copies/mL (Figure. 6B). GE ranges from 227.5 to 53047.5 copies, with the median being 1143.75 copies, were used as input for the pre-amplification reaction (Figure. 6C). Taking 0.25% as an average LOD assay cutoff, a minimum of 500 total copies would be required as input to be able to detect down to 0.25% allele burden. Indeed, in a small set of samples with sample input GE numbers ranging from 300-1500, the detected variant allele frequency (VAFs) ranged from 0.2% to 1% with our in-house assay. Based on these values, the calculated mutant copy numbers that we were able to detect were in a single digit range (Supplementary Table 6). This suggests that for each sample, the LOD is determined by not only the assay LOD, but also how many mutant copies existed in the input material (sample driven LOD). For samples that have <500 GEs, increasing the pre-amplification reaction volume would be required to raise the sample-driven LOD.

To understand the accuracy of the in-house MX-PCR-ddPCR assays, concordance with the Inostics assay was assessed using a total of 38 positive samples, with mutant allele frequencies ranging from 68.4% to 0.059%. A good concordance (R^2^=0.9871, p<0.0001) was observed (Figure 6D) between the two platforms. Only two data points with allele frequencies of 0.06 measured in Inostics assay could not be detected by the in-house assays. Eight assays that were tested negative by BEAMing assay were also negative in our assay.

The reasoning behind the use of a pre-amplification step was to save precious samples for future application and testing. But this introduced two concerns. First, pre-amplification did result in higher background, thus driving up the LOD/LOL of assays. Second, there was a risk that pre-amplification could compromise the quantitative nature of the assay. To address these concerns, two cfDNA samples previously characterized by BEAMing were used to compare results with or without pre-amplification using our method. One of them had extremely high GE copies (980,577 copies/mL plasma) and possessed PIK3CA E542K mutation at 0.059% allele burden. The other patient sample had lower GEs (15093 copies/mL plasma), but contained multiple mutations with relatively higher allele frequencies, such as 2.6% of *ESR1* Y537N, 29.8% of ESR1 Y537C, 0.925% of ESR1 E380Q and 51.1% of *PIK3CA* E545K. When allele frequencies of mutations with and without multiplex amplification were measured, comparable results were observed (Table 2). Again, the only sample that had inconsistent result was from the sample that had a 0.059% allele frequency, which is below the LOD of the assay. This result further confirms the maintenance of the quantitative nature of the assay after multiplex pre-amplification.

**Table 2.** VAF measured in patient samples with and without pre-amplification, and concordence to BEAMing assays. “-” means not measured.

| **Assay** | **Patient 1**<br>**(BEAMing)** | **Patient 1**<br>**without pre-**<br>**amp** | **Patient 1 with**<br>**pre-amp** | **Patient 2**<br>**(BEAMing)** | **Patient 2 without**<br>**pre-amp** | **Patient 2**<br>**with pre-amp** |
| --- | --- | --- | --- | --- | --- | --- |
| **cfDNA Input**<br>**amount** | **1160991 GE** | **33525 GE** | **125720 GE** | **31239 GE** | **1295 GE** | **1942 GE** |
| ESR1 Y537N | Neg | Neg | Neg | 2.6 (Pos) | 4.6 (Pos) | 4.02 (Pos) |
| ESR1 Y537C | Neg | Neg | Neg | 29.8 (Pos) | 35.4 (Pos) | 35.2 (Pos) |
| ESR1 S463P | Neg | Neg | - | Neg | Neg | - |
| ESR1 E380Q | Neg | Neg | Neg | 0.925 (Pos) | 1.1 (Pos) | 1.5 (Pos) |
| PIK3CA<br>C420R | Neg | Neg | - | Neg | Neg | - |
| PIK3CA<br>E542K | 0.059 (Pos) | 0.12 (Pos) | 0.128 (False<br>Neg) | Neg | 0.29 (False Pos) | Neg |
| PIK3CA<br>E545K | Neg | Neg | Neg | 51.1 (Pos) | 55.6 (Pos) | 56.3 (Pos) |
| PIK3CA<br>M1043I | Neg | Neg | - | Neg | Neg | - |

## Discussion

In this paper, we validated a panel of ddPCR assays for detecting *ESR1* and *PIK3CA* mutations following MX-PCR amplification. The motivation to design and validate such a panel was to develop a sensitive, reproducible and cost-effective way to longitudinally track clinically relevant *ESR1* and *PIK3CA* mutations in MBC using a user-friendly and de-centralized workflow. The potential of tracking PI3K and *ESR1* mutation dynamics in plasma as biomarkers to predict clinical response to endocrine therapy has been the focus of many clinical studies, such as BOLERO-2 phase 3 study ^7,39^, Paloma-3 Phase III Study ^4^, and the AZD9496 Phase I study ^5^, and development of custom panels like the one described here is critical to obtain reliable cfDNA mutation assessment while using as little material as possible.

ddPCR has several advantages compared to NGS methods for detecting and monitoring cfDNA mutations. First, it offers higher sensitivity than many off-the-shelf NGS panels. The advantage of NGS panels is simultaneous detection of hotspot and de-novo mutations which is ideal for discovery applications; however, the vast content often results in lower sensitivity. Breakthroughs in noise reduction from NGS by using molecular indices ^17,29,40^ have partially addressed this limitation with many commercially available panels achieving LODs below 1% ^8^. For example, Archer Reveal’s detection rate is 1% with 10ng of cfDNA (could be lower with more input cfDNA) ^34^and the FACT assay requirement >20 ng of cfDNA reportedly achieves a 0.5% detection limit ^33^. While these assays are suitable for baseline characterization of patients and treatment selection, quantitatively measuring allele frequencies below 1% after treatment longitudinally is still challenging and cost-prohibitive. Several studies have shown a high degree of variability among commercial assays, particularly at low allele fractions (AF<1%) ^36,41^, suggesting that the commercially available assays may not be ideal for monitoring low allele frequencies like that observed in ESR1. NGS assays also come with a high sample amount requirement and high resource requirement from both the wet-lab and informatics component of the assay. In comparison, ddPCR assays while restricting detection to specific mutations allows for higher sensitivity which is advantageous in clinical applications such as monitoring response to therapy and detection of relapse. O’Leary and colleagues demonstrated that rare mutations of *ESR1* and *PIK3CA* existing at baseline that could be identified by ddPCR assays were not identified by targeted NGS panels ^3^. These mutations were all below 1%. Furthermore, the high sensitivity, absolute quantification nature and simplified analysis of the ddPCR technology make it a clinic-friendly approach. A number of reports have highlighted its superior accuracy ^42^. Rapid microfluidic analysis of thousands of droplets per sample makes ddPCR practical for routine use ^43,44^. Indeed, we were able to validate the LOL to 0.25% for most of the assays after MX-PCR amplification, while LOD was even lower at 0.1%.

The second advantage of this method is the flexibility of pre-amplification. After MX-PCR amplification, each mutation can be measured independently. Therefore, one can choose to measure all the mutations in a custom panel or measure selected mutations resulting in cost flexibility. The third advantage is that pre-amplification lowers the assay input requirement which can be limiting in clinical samples allowing for banking cfDNA for future applications. With the use of MX-PCR amplification, <10% of a patient’s cfDNA sample, that is derived typically from 2-3 mL of plasma, gets used. In contrast, based on our own calculation, the majority of samples would be exhausted after mutation tracking using 4-5 regular singleplex ddPCR assays.

During the course of assay development and validation, one significant challenge was to identify the appropriate sample type and matrix for control samples. Patient-derived plasma samples have a very limited amount of cfDNA and were expensive to purchase, limiting their utility as sample of choice for control material. Additionally, as up to 20 assays needed to be validated, it was very hard and almost impossible to find human cfDNA samples that contained all the mutations of interest. Hence contrived samples were the only viable choice for assay development and validations. The two most commonly used contrived samples are cell line gDNA spiked in with mutant gBlocks or a mixture of wildtype (WT) and mutant gBlocks. We tried the former at the beginning to preserve the heterogeneous nature of gDNA as opposed to specific amplicon based gBlocks. We had to switch to a gBlock only solution, since it required too much cell line gDNA to generate samples for LOD and LOL measurement. For example, to generate samples that contained 0.05% of mutant allele frequencies, at least 5 copies of mutant DNA were needed to be spiked into 10^4^ copies of WT DNA. To minimize random sampling error, 50 copies of mutant DNA were spiked into 10^5^ copies of WT DNA. To complete validation for all 21 assays including multiple conditions (with and without pre-amplification), the amount of gDNA required was over 5mg and became unfeasible. In addition, each batch of gDNA was needed to be characterized for copy numbers and alterations to ensure the base truth of the samples. Hence we switched to the more reproducible and cost-effective gBlocks. But they have their own limitations since gBlocks are very homogeneous, hence raising the concern that this sample type would generate an artificially higher background than cell line gDNA or patient derived cfDNA, which contain greater sequence diversity. Indeed, when the background was compared between gBlocks and cfDNA references samples, it was noticed that gBlocks had higher background than cfDNA references and patient cfDNAs for many assays. However, for other assays, background of gBlock and cfDNA references remained comparable. Overall our studies and data indicate the importance and downstream implications of choice of sample type for use in assay development and validation, and it could be the difference between a positive or a false negative call from the assay.

In summary, we have developed a panel of ddPCR assays following MX-PCR amplification for detecting *ESR1* and *PIK3CA* mutations. The method has demonstrated good sensitivity, reproducibility and accuracy. As applications of cfDNA expand, we expect a growing number of clinical correlative studies to understand the value of cfDNA modulation as surrogate readouts of efficacy and response, further highlighting the need for custom panels like ours.

## Material and Methods

### Samples

Wild type and mutant gBlocks used in the validation were ordered from Integrated DNA Technologies (IDT). Their sequences were listed in Supplementary Table 7. Multiplex HD780 cfDNA Reference Standard Set were ordered from Horizon. Ninety plasma samples were collected through an ongoing clinical trial sponsored by H3 Biomedicine: Trial of H3B-6545, in Women with Locally Advanced or Metastatic Estrogen Receptor-positive, HER2 Negative Breast Cancer (NCT03250676). An institutional review board or equivalent approved the study at each participating site, with patients supplying written informed consent.

### Multiplex PCR

Multiplex PCR primers were designed using the Primer3 website (http://bioinfo.ut.ee/primer3-0.4.0/) and checked for specificity using the NCI BLAST Standard Nucleotide website (https://blast.ncbi.nlm.nih.gov/Blast.cgi). Multiplex PCR primers sequences were listed in Supplementary Table 2. The primers reconstituted according to IDT instructions into stock solutions at 100μM with TE buffer (Low-EDTA). NanoDrop is used to check the OD260 of the primers stock solution. OD260 was measured in triplicate and average was used for calculation of primer concentration. The ratio of nmol/OD260 used were as following: ESR1m-538-1F (5.43 nmol/OD260), ESR1m-538-1R (6.38), ESR1m-463-F (4.81), ESR1m-463-R (4.63), ESR1m-380-F (4.78), ESR1m-380-R (5.28), PIK3CAm-420-F (4.05), PIK3CAm-420-R (4.99), PIK3CAm-545-F (3.22), PIK3CAm-545-R (3.5), PIK3CAm-1047-1F (4.26), PIK3CAm-1047-1R (4.96). The forward and reverse primers for each amplicon is mixed separately in separate tubes (six tubes in total) to obtain the final concentration of each primer 10uM.

To optimize the multiplex PCR primer ratio, the MX-PCR was performed with a total reaction volume of 25 μL containing 12.5 μL Phusion Hot Start Flex 2X Master Mix (New England BioLabs), 25 ng of K562 cell line genomic DNA and 5 μL of the primer pool as defined in Supplementary Table 3. Amplification was carried out on a C1000 Touch PCR System (Bio-Rad) using the following conditions: an initial denaturation step at 98 °C for 30 sec, 15 cycles of 98 °C for 10 sec, 66 °C for 20 sec, decreasing 0.5 °C per cycle, and 72 °C for 20 sec, followed by 10 or 15 cycles of 98 °C for 10 sec, 59 °C for 20 sec, 72 °C for 20 sec and a final extension step of 72 °C for 5 min. For final production, 15 cycles of final extension was selected.

### Sanger sequencing

After PCR products were purified with Agencourt^®^ AMPure XP beads (Beckman Coulter), the cycle sequencing reactions were carried out using standard Sanger sequencing methods with BigDye v3.1 (Thermo Fisher Scientific) on a C1000 Touch PCR System (Bio-Rad). The excess dye-terminators in the reactions were removed with Agencourt CleanSEQ^®^ paramagnetic beads (Beckman Coulter). Bi-directional sequencing was performed on an ABI 3730XL DNA Analyzer (Life Technologies). Sequencher™ 5.2.3 software (Gene Codes Corp) was used for post-sequence analysis.

### Sequencing by Ion Torrent Platform

Ion Plus Fragment libraries were prepared for the MX-PCR products using the Ion Plus Fragment Library Kit (Thermo Fisher Scientific). The barcoded libraries were then quantified, normalized and pooled at a concentration of 50 pM. Template preparation was performed using the Ion PGM™ Template OT2 200 Kit (Thermo Fisher Scientific) on the Ion OneTouch™ 2 System (Thermo Fisher Scientific). The templates were sequenced on the Ion PGM™ System, using the Ion PGM™ sequencing 200 Kit v2 and the Ion 314™ Chip Kit v2 (Thermo Fisher Scientific). Data from the PGM runs were processed using the Ion Torrent Suite 5.4 software to generate sequencing reads. After sequence alignment, the amplicon coverage report was used for further analysis. The total reads, which represent the total number of targeted PCR products for each amplicon at each of the primer ratios, were compared.

### cfDNA extraction and quantification

Blood was collected by standard phlebotomy techniques in Cell-Free DNA BCT tubes (Streck, La Vista, NE; referred to as cfDNA BCTs). All tubes were filled to 10 mL as recommended by the manufacturer. After the blood draw, tubes were transported at room temperature (RT) to Sysmex Inostics’ (Baltimore, MD) laboratory for processing and storage. For plasma preparation, blood tubes were centrifuged at 1,600 × g for 10 min at RT using a swing-out rotor. To remove any residual blood cells, the supernatant was centrifuged a second time at 6,000 × g for 10 min at RT using a fixed-angle rotor and a smooth breaking profile. cfDNA was extracted from 2 mL of plasma at baseline and from 3 mL of plasma post-treatment using QIAamp DNA purification kit (Qiagen, Venlo, the Netherlands) according to the manufacturer’s instructions. The total amount of amplifiable human genomic DNA purified from plasma samples was quantified using a modified version of human long interspersed element 1 (LINE-1) real-time PCR assay and reported as genome equivalents (GE), with one GE being one haploid human genome weighing 3.3 pg ^13,45^.

### gBlocks preparation for validation

Wild type and mutant gBlocks controls from IDT (sequence in the table before) were reconstituted into stock solutions at ~10 ng/µL with TE buffer (Low-EDTA). NanoDrop checked concentration was used to make a stock of 1 ng/uL. The total gBlock copies in 1 ng of the gBlocks were calculated using the following formula:

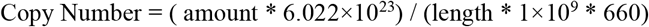

First, mutant and wild type were diluted to 10^5^ copies/μL. To generate standard curve, the following VAF 50%, 25%, 10%, 5%, 1%, 0.5%, 0.25%, 0.1%, 0.05%, 0.01% were prepared for individual mutant gBlocks.

### ddPCR setup

After pre-amplification, samples were diluted 10^6^ times with nuclease-free water. The first run is set up to quick check of copy number in each sample for each assay. The target concentration for the first run was 1500-3500 copies/uL. If not in the range, concentration should be adjusted for the next plate. The second plate is to determine the real VAF per assay. Every sample was measured in triplicate. A no-DNA water control and a positive control sample (ex. relevant pre-clinical samples or GBlocks without pre-amp) were included for each assay. All ddPCR reactions were tested using Bio-Rad QX200™ Droplet Digital™ PCR System with automated droplet generator, following Bio-Rad instruction (Bio-Rad laboratories, CA). The assays used are shown in Table 1. ddPCR were performed in a 22 μL reaction containing 11 μL ddPCR™ Supermix for Probes (No dUTP), 1.1 μL of assay and up to 9.9 uL of sample. ddPCR condition for assays is: 95°C for 10 min, followed by 40 cycles of 94°C for 30 sec, assay specific Tm for 60 sec (Ramp rate 2°C/sec), and final incubation 98°C for 10 min. Assay specific Tm are shown in Table 1.

### Data analysis

The ddPCR data were analyzed with QuantaSoft analysis software version 1.7 (Bio-Rad). The positive droplets containing amplified products were discriminated from negative droplets by applying a threshold above the negative droplets. Each assay is gated according to positive and negative controls using 2D Amplitude data.

### Statistical analysis

Linear regression for analyzing the limited of linearity of the ddPCR assay was analyzed with GraphPad Prism v7.02.

### Data availability

The datasets generated during and/or analyzed during the current study are available from the corresponding author on reasonable request.

## Contributions

Study concept and design: H.Y., P.K., G.H. acquisition of data: H.Y., S.D., C.K., G.H, H.G., S.F., A.R. analysis and interpretation of data: H.Y., Z.Z., P.K., V.R. Draft the manuscript: H.Y., C.K., Z.Z., V.R. critical revision of the manuscript for important intellectual content: M.K., J.S., P.G.S., L.Y., P.Z., P.K.

## References

1. Ghoncheh, M., Pournamdar, Z. & Salehiniya, H. Incidence and Mortality and Epidemiology of Breast Cancer in the World. Asian Pacific journal of cancer prevention: APJCP 17, 43–6 (2016).

2. Effects of chemotherapy and hormonal therapy for early breast cancer on recurrence and 15-year survival: an overview of the randomised trials. Lancet 365, 1687–717 (2005).

3. O’Leary, B. et al. The Genetic Landscape and Clonal Evolution of Breast Cancer Resistance to Palbociclib plus Fulvestrant in the PALOMA-3 Trial. Cancer discovery 8, 1390–1403 (2018).

4. O’Leary, B. et al. Early circulating tumor DNA dynamics and clonal selection with palbociclib and fulvestrant for breast cancer. Nature communications 9, 896 (2018).

5. Paoletti, C. et al. Circulating Biomarkers and Resistance to Endocrine Therapy in Metastatic Breast Cancers: Correlative Results from AZD9496 Oral SERD Phase I Trial. Clinical cancer research: an official journal of the American Association for Cancer Research 24, 5860–5872 (2018).

6. Fribbens, C. et al. Tracking evolution of aromatase inhibitor resistance with circulating tumour DNA analysis in metastatic breast cancer. Annals of oncology: official journal of the European Society for Medical Oncology 29, 145–153 (2018).

7. Chandarlapaty, S. et al. Prevalence of *ESR1* Mutations in Cell-Free DNA and Outcomes in Metastatic Breast Cancer: A Secondary Analysis of the BOLERO-2 Clinical Trial. JAMA oncology 2, 1310–1315 (2016).

8. Masunaga, N. et al. Highly sensitive detection of *ESR1* mutations in cell-free DNA from patients with metastatic breast cancer using molecular barcode sequencing. Breast cancer research and treatment 167, 49–58 (2018).

9. Fribbens, C. et al. Plasma *ESR1* Mutations and the Treatment of Estrogen Receptor-Positive Advanced Breast Cancer. Journal of clinical oncology: official journal of the American Society of Clinical Oncology 34, 2961–8 (2016).

10. Clatot, F. et al. Kinetics, prognostic and predictive values of *ESR1* circulating mutations in metastatic breast cancer patients progressing on aromatase inhibitor. Oncotarget 7, 74448–74459 (2016).

11. Miller, T.W., Balko, J.M. & Arteaga, C.L. Phosphatidylinositol 3-kinase and antiestrogen resistance in breast cancer. Journal of clinical oncology: official journal of the American Society of Clinical Oncology 29, 4452–61 (2011).

12. Spoerke, J.M. et al. Heterogeneity and clinical significance of *ESR1* mutations in ER-positive metastatic breast cancer patients receiving fulvestrant. Nature communications 7, 11579 (2016).

13. Diehl, F. et al. Circulating mutant DNA to assess tumor dynamics. Nature medicine 14, 985–90 (2008).

14. Dawson, S.J., Rosenfeld, N. & Caldas, C. Circulating tumor DNA to monitor metastatic breast cancer. The New England journal of medicine 369, 93–4 (2013).

15. Wan, J.C.M. et al. Liquid biopsies come of age: towards implementation of circulating tumour DNA. Nature reviews. Cancer 17, 223–238 (2017).

16. Yao, W., Mei, C., Nan, X. & Hui, L. Evaluation and comparison of in vitro degradation kinetics of DNA in serum, urine and saliva: A qualitative study. Gene 590, 142–8 (2016).

17. Dawson, S.J. et al. Analysis of circulating tumor DNA to monitor metastatic breast cancer. The New England journal of medicine 368, 1199–209 (2013).

18. Marchetti, A. et al. Early Prediction of Response to Tyrosine Kinase Inhibitors by Quantification of EGFR Mutations in Plasma of NSCLC Patients. Journal of thoracic oncology: official publication of the International Association for the Study of Lung Cancer 10, 1437–43 (2015).

19. Tie, J. et al. Circulating tumor DNA as an early marker of therapeutic response in patients with metastatic colorectal cancer. Annals of oncology: official journal of the European Society for Medical Oncology 26, 1715–22 (2015).

20. Garlan, F. et al. Early Evaluation of Circulating Tumor DNA as Marker of Therapeutic Efficacy in Metastatic Colorectal Cancer Patients (PLACOL Study). Clinical cancer research: an official journal of the American Association for Cancer Research 23, 5416–5425 (2017).

21. Gray, E.S. et al. Circulating tumor DNA to monitor treatment response and detect acquired resistance in patients with metastatic melanoma. Oncotarget 6, 42008–18 (2015).

22. Schreuer, M. et al. Quantitative assessment of BRAF V600 mutant circulating cell-free tumor DNA as a tool for therapeutic monitoring in metastatic melanoma patients treated with BRAF/MEK inhibitors. Journal of translational medicine 14, 95 (2016).

23. Sorenson, G.D. et al. Soluble normal and mutated DNA sequences from single-copy genes in human blood. Cancer epidemiology, biomarkers & prevention: a publication of the American Association for Cancer Research, cosponsored by the American Society of Preventive Oncology 3, 67–71 (1994).

24. Ou, S.I., Nagasaka, M. & Zhu, V.W. Liquid Biopsy to Identify Actionable Genomic Alterations. American Society of Clinical Oncology educational book. American Society of Clinical Oncology. Annual Meeting, 978–997 (2018).

25. Vogelstein, B. & Kinzler, K.W. Digital PCR. Proceedings of the National Academy of Sciences of the United States of America 96, 9236–41 (1999).

26. Diehl, F. et al. BEAMing: single-molecule PCR on microparticles in water-in-oil emulsions. Nature methods 3, 551–9 (2006).

27. Forshew, T. et al. Noninvasive identification and monitoring of cancer mutations by targeted deep sequencing of plasma DNA. Science translational medicine 4, 136ra68 (2012).

28. Gale, D. et al. Analytical performance and validation of an enhanced TAm-Seq circulating tumor DNA sequencing assay. in the 107th Annual Meeting of the American Association for Cancer Research Vol. 76 (Cancer Research, New Orleans, LA. Philadelphia (PA), 2016).

29. Kinde, I., Wu, J., Papadopoulos, N., Kinzler, K.W. & Vogelstein, B. Detection and quantification of rare mutations with massively parallel sequencing. Proceedings of the National Academy of Sciences of the United States of America 108, 9530–5 (2011).

30. Newman, A.M. et al. An ultrasensitive method for quantitating circulating tumor DNA with broad patient coverage. Nature medicine 20, 548–54 (2014).

31. Lanman, R.B. et al. Analytical and Clinical Validation of a Digital Sequencing Panel for Quantitative, Highly Accurate Evaluation of Cell-Free Circulating Tumor DNA. PloS one 10, e0140712 (2015).

32. Schwaederle, M. et al. Detection rate of actionable mutations in diverse cancers using a biopsy-free (blood) circulating tumor cell DNA assay. Oncotarget 7, 9707–17 (2016).

33. Clark, T.A. et al. Analytical Validation of a Hybrid Capture-Based Next-Generation Sequencing Clinical Assay for Genomic Profiling of Cell-Free Circulating Tumor DNA. The Journal of molecular diagnostics: JMD 20, 686–702 (2018).

34. Lee, J.E. et al. Anchored Multiplex PCR Enables Sensitive and Specific Detection of Variants in Circulating Tumor DNA by Next-Generation Sequencing. in Molecular Med Tri-Con (San Francisco, CA, 2017).

35. Page, K. et al. Next Generation Sequencing of Circulating Cell-Free DNA for Evaluating Mutations and Gene Amplification in Metastatic Breast Cancer. Clinical chemistry 63, 532–541 (2017).

36. Stetson, D. et al. Examination of analytical factors impacting concordance of plasma-tumor testing by next-generation sequencing (NGS). in American Association for Cancer Research Annual Meeting (Cancer Research, Washington, DC. Philadelphia (PA), 2017).

37. Stetson, D. et al. Examination of intra-assay variability in a commercial ctDNA NGS assay with triplicate clinical plasma and pooled normal samples in American Association for Cancer Research Annual Meeting 2018 (Cancer Research, Chicago, IL. Philadelphia (PA), 2018).

38. Takeshita, T. et al. Clinical significance of monitoring *ESR1* mutations in circulating cell-free DNA in estrogen receptor positive breast cancer patients. Oncotarget 7, 32504–18 (2016).

39. Moynahan, M.E. et al. Correlation between *PIK3CA* mutations in cell-free DNA and everolimus efficacy in HR(+), HER2(-) advanced breast cancer: results from BOLERO-2. British journal of cancer 116, 726–730 (2017).

40. Kivioja, T. et al. Counting absolute numbers of molecules using unique molecular identifiers. Nature methods 9, 72–4 (2011).

41. Torga, G. & Pienta, K.J. Patient-Paired Sample Congruence Between 2 Commercial Liquid Biopsy Tests. JAMA oncology 4, 868–870 (2018).

42. Hindson, C.M. et al. Absolute quantification by droplet digital PCR versus analog real-time PCR. Nature methods 10, 1003–5 (2013).

43. Hindson, B.J. et al. High-throughput droplet digital PCR system for absolute quantitation of DNA copy number. Analytical chemistry 83, 8604–10 (2011).

44. Pinheiro, L.B. et al. Evaluation of a droplet digital polymerase chain reaction format for DNA copy number quantification. Analytical chemistry 84, 1003–11 (2012).

45. Rago, C. et al. Serial assessment of human tumor burdens in mice by the analysis of circulating DNA. Cancer research 67, 9364–70 (2007).

