## supplementary tables for "Coupling multiplex pre-amplification and droplet digital PCR for longitudinal monitoring of *ESR1* and *PIK3CA* mutations from plasma cell-free DNA"

**Supplementary Table 1. Overview of the panel design.** Individual mutation frequency in metastatic breast cancer and previous used methods are shown.

| Basic Panel Information | | | | | Published panel and methods | | | | | |
|---|---|---|---|---|---|---|---|---|---|---|
| Amplicon | Gene | AA Change | cDNA Change | Cosmic ID | Clatot F et al, Oncotarget, 2016 | Paoletti C et al., Clin Cancer Res, 2018, | Chandarlapaty S et al, JAMA Oncol, 2016; Moynahan, M.E. BJC, 2017 | Takeshita T et al., Oncotarget, 2017 | Spoker JM et al., Nat Commun, 2016 | O'Leary B et al., Nat Commun, 2018 |
| 1 | ESR1 | V534E | c.1601T>A | COSM4774827 | Not measured | multiplex ddPCR | Not measured | Not measured | OncoBEAMTM CLIA Panel 1 | Not measured |
| | | P535H | c.1604C>A | COSM4944018 | Not measured | Not measured | Not measured | Not measured | OncoBEAMTM CLIA Panel 1 | Not measured |
| | | L536H | c.1607T>A | N/A | Not measured | Not measured | Not measured | Not measured | OncoBEAMTM CLIA Panel 1 | Not measured |
| | | L536R | c.1607T>G | COSM4774826 | Not measured | multiplex ddPCR | Not measured | Not measured | OncoBEAMTM CLIA Panel 1 | Biorad multiplex1 |
| | | L536Q | c.1607_1608TC>AG | N/A | Not measured | multiplex ddPCR | Not measured | Not measured | OncoBEAMTM CLIA Panel 1 | Not measured |
| | | Y537S | c.1610A>C | COSM1074639 | singleplex ddPCR | singleplex ddPCR | singleplex ddPCR | singleplex ddPCR, LBx ESR1 screen | OncoBEAMTM CLIA Panel 1 | Biorad multiplex2 |
| | | Y537N | c.1609T>A | COSM1074635 | singleplex ddPCR | singleplex ddPCR | Not measured | singleplex ddPCR, LBx ESR1 screen | OncoBEAMTM CLIA Panel 1 | Biorad multiplex2 |
| | | Y537C | c.1610A>G | COSM1074637 | singleplex ddPCR | singleplex ddPCR | Not measured | singleplex ddPCR | OncoBEAMTM CLIA Panel 1 | Biorad multiplex1 |
| | | D538G | c.1613A>G | COSM94250 | singleplex ddPCR | singleplex ddPCR | singleplex ddPCR | singleplex ddPCR, LBx ESR1 screen | OncoBEAMTM CLIA Panel 1 | Biorad multiplex1 |
| 2 | ESR1 | S463P | c.1387T>C | COSM4771561 | Not measured | Not measured | Not measured | Not measured | OncoBEAMTM CLIA Panel 1 | Biorad multiplex2 |
| 3 | ESR1 | E380Q | c.1138G>C | COSM3829320 | Not measured | singleplex ddPCR | Not measured | Not measured | OncoBEAMTM CLIA Panel 1 | Biorad multiplex1 |

| | | | | | | | | | | |
|---|---|---|---|---|---|---|---|---|---|---|
| 4 | PIK3 CA | C420 R | c.1258T>C | COSM757 | Not measured | Not measured | Not measured | Not measured | OncoBEAMTM CLIA Panel 1 | Not measured |
| 5 | PIK3 CA | E542 K | c.1624G>A | COSM760 | Not measured | Not measured | singleplex ddPCR | LBx PIK3CA screen 1 | OncoBEAMTM CLIA Panel 1 | singleplex ddPCR |
| | | E545 G | c.1634A>G | COSM764 | Not measured | Not measured | Not measured | LBx PIK3CA screen 1 | OncoBEAMTM CLIA Panel 1 | Not measured |
| | | E545 K | c.1633G>A | COSM763 | Not measured | Not measured | singleplex ddPCR | LBx PIK3CA screen 1 | OncoBEAMTM CLIA Panel 1 | singleplex ddPCR |
| | | Q546 K | c.1636C>A | COSM766 | Not measured | Not measured | Not measured | LBx PIK3CA screen 1 | OncoBEAMTM CLIA Panel 1 | Not measured |
| 6 | PIK3 CA | M104 3I | c.3129G>A | COSM2931 3 | Not measured | Not measured | Not measured | LBx PIK3CA screen 2 | OncoBEAMTM CLIA Panel 1 | Not measured |
| | | M104 3I | c.3129G>T | COSM773 | Not measured | Not measured | Not measured | LBx PIK3CA screen 2 | OncoBEAMTM CLIA Panel 1 | Not measured |
| | | H1047 L | c.3140A>T | COSM776 | Not measured | Not measured | Not measured | LBx PIK3CA screen 2 | OncoBEAMTM CLIA Panel 1 | singleplex ddPCR |
| | | H1047 R | c.3140A>G | COSM775 | Not measured | Not measured | singleplex ddPCR | LBx PIK3CA screen 2 | OncoBEAMTM CLIA Panel 1 | singleplex ddPCR |
| | | H1047 Y | c.3139C>T | COSM774 | Not measured | Not measured | Not measured | LBx PIK3CA screen 2 | OncoBEAMTM CLIA Panel 1 | Not measured |

**Supplementary Table 2. The prevalence of ESR1 and PIK3CA mutations among ER+ HER2- metastatic breast cancer patients.** “-“means data not available.

| Source | ESR1 mutations | PIK3CA mutations | ESR1 and/or PIK3CA mutations | Total # of pts |
|---|---|---|---|---|
| Toy W. et al., 2013 | 25.0% | - | - | 36 |
| Robinson D. et al., 2013 | 54.5% | - | - | 11 |
| Marenbach, K. et al., 2013 | 38.5% | - | - | 13 |
| Jeselsohn, R. et al., 2014 | 14.5% | - | - | 76 |
| Chandarlapaty, S. et al., 2016 | 28.8% | - | - | 156 |
| Fumagalli D. et al., 2016 | 10.8% | 33.3% | - | 87 |
| Spoerke JM. Et al., 2016 | 37.3% | 40.5% | 55.6% | 153 |

**Supplementary Table 3. Primer sequences for the targeted amplicons.**

| Amplicon Name | Primer Name | Sequence (5'- 3') | PCR Size (bp) |
|---|---|---|---|
| E380 | ESR1m-380-F | TTCATTTGAGTCAGCAGGGTTT | 157 |
| | ESR1m-380-R | GCAAACAGTAGCTTCCCTGG | |
| E463 | ESR1m-463-F | TGAACACTCTGGGTCTCCTAGA | 170 |
| | ESR1m-463-R | TCAGGTGGATCAAAGTGTCTGT | |
| E538 | ESR1m-538-1F | AGTCCTTTCTGTGTCTTCCCA | 173 |
| | ESR1m-538-1R | GCTTTGGTCCGTCTCCTCC | |
| P420 | PIK3CAm-420-F | GAGATGATTGTTGAATTTTCCTTTTG | 186 |
| | PIK3CAm-420-R | ACTGGCCAAAGATTCAAAGC | |
| P545 | PIK3CAm-545-F | CTAGAGACAATGAATTAAGGGAAAATGAC | 134 |
| | PIK3CAm-545-R | GAATCTCCATTTTAGCACTTACCTGTGA*C*T*C | |
| P1047 | PIK3CAm-1047-1F | TTTGATGACATTGCATACATTCGA | 165 |
| | PIK3CAm-1047-1R | TTTTCAGTTCAATGCATGCTGT | |

* Phosphorothioate bond modification

**Supplementary Table 4. Primer ratios tested to determine optimal amplification of all targets.** Each pair of primers was first diluted to 10µM. Different pool of primers were then generated by combining the six pairs of primers at different ratios, showing below:

| | | Amplicon | | | | | |
|---|---|---|---|---|---|---|---|
| | | E538 | E463 | E380 | P420 | P545 | P1047 |
| Primer Pool | A | 1 | 1 | 1 | 1 | 1 | 1 |
| | B | 1 | 1 | 1 | 2 | 1 | 1.5 |
| | C | 1 | 1 | 1 | 3 | 1 | 1.5 |
| | D | 0.5 | 1 | 1 | 4 | 1.5 | 1 |
| | E | 0.5 | 0.75 | 1 | 4 | 2 | 1 |
| | F | 0.5 | 1 | 1 | 3 | 1.5 | 0.75 |
| | G | 0.75 | 1 | 1 | 4 | 1.5 | 1 |
| | H | 0.75 | 0.75 | 1 | 4 | 2 | 1 |
| | I | 0.75 | 1 | 1 | 3 | 1.5 | 0.75 |

**Supplementary Table 5. Reproducibility of the assays using contrived gBlock samples.** Average measured VAF (%) and corresponding CV% are shown for representative assays. Three independent experiments from preamplification to ddPCR measurement are shown here.

| ESR1 L536H | | ESR1 S463P | | ESR1 E380Q | | PIK3CA C420R | | PIK3CA H1047R | |
|---|---|---|---|---|---|---|---|---|---|
| Average Measured (%) | CV% | Average Measured (%) | CV% | Average Measured (%) | CV% | Average Measured (%) | CV% | Average Measured (%) | CV% |
| 5.94 | 9.13 | 6.48 | 13.16 | 6.57 | 3.94 | 6.70 | 1.13 | 2.98 | 4.14 |
| 2.49 | 13.32 | 2.15 | 23.32 | 2.07 | 6.03 | 2.14 | 15.03 | 0.76 | 11.87 |
| 0.38 | 33.33 | 0.34 | 12.62 | 0.30 | 61.56 | 0.31 | 12.17 | 0.14 | 26.57 |
| 0.21 | 29.06 | 0.31 | 15.55 | 0.21 | 38.08 | 0.24 | 27.63 | 0.13 | 3.02 |
| 0.15 | 31.89 | 0.20 | 36.61 | 0.06 | 54.85 | 0.18 | 29.97 | 0.10 | 34.90 |

**Supplementary Table 6. VAF% and GE numbers in selected patient samples that had low GE number and low VAF%.**

| ddPCR | Inostics VAF (%) | In-house VAF (%) | In-house total GE (based on LINE-1 quantification) | Calculated mutant copies in the input (Based on H3 VAF) |
|---|---|---|---|---|
| ESR1 D538G | 0.15 | 0.27 | 502.5 | 1.36675 |
| ESR1 D538G | 0.14 | 0.21 | 1335 | 2.83 |
| ESR1 D538G | 0.82 | 0.32 | 1177.5 | 3.77 |
| PIK3CA H1047R | 0.56 | 1.24 | 307.5 | 3.81 |
| ESR1 L536H | 0.17 | 0.17 | 2880 | 4.90 |
| ESR1 Y537N | 0.32 | 0.59 | 1177.5 | 6.95 |

**Supplementary Table 7. gBlock sequences used in panel validation.**
See excel file.

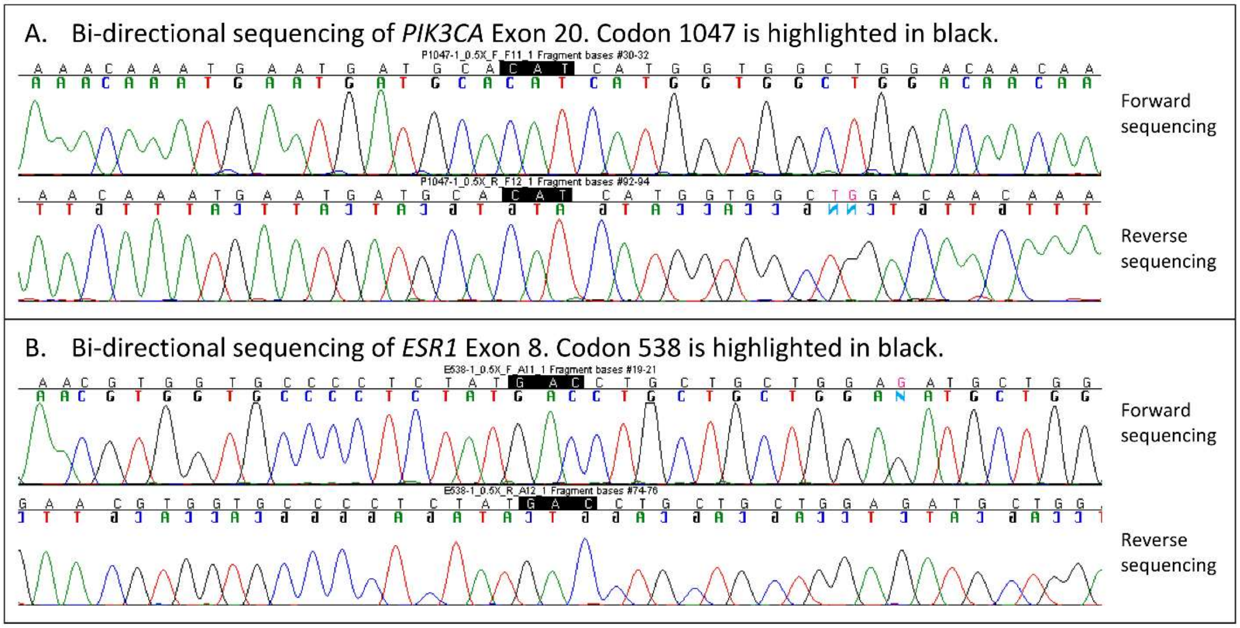


**Supplementary Figure 1.** Examples of the Sanger sequencing of (A) *PIK3CA* Exon 20 codon 1047 and (B) *ESR1* exon 8 codon 538.
